## Supplementary material for "Paramyxoviruses in Old World fruit bats (Pteropodidae): an open database and synthesis of sampling effort, viral positivity, and coevolution": Table S1

**Supplementary Table 1. Database breakdown by sample type and detention method.** Sample size (records and number of samples), percentage of records with at least one positive sample, and average prevalence by detection method, sample type, and viral subfamily.

| | All PMVs | | | | Orthoparamyxovirinae | | | | Rubulavirinae | | | |
|---|---|---|---|---|---|---|---|---|---|---|---|---|
| Sample type | *n* rows | % positive rows | *n* samples | Average % prevalence | *n* rows | % positive rows | *n* samples | Average % prevalence | *n* rows | % positive rows | *n* samples | Average % prevalence |
| **Serology** | | | | | | | | | | | | |
| Blood or serum | 395 | 74.73 | 31675 | 25.69 | 327 | 76.38 | 28832 | 26.53 | 68 | 66.67 | 2843 | 21.55 |
| **PCR** | | | | | | | | | | | | |
| Blood or serum | 43 | 16.28 | 4867 | 0.53 | 42 | 16.67 | 4828 | 0.55 | 0 | - | 0 | - |
| Fecal, rectal, anal | 54 | 14.81 | 2546 | 0.78 | 19 | 42.11 | 2326 | 2.22 | 0 | - | 0 | - |
| Feces (single) | 17 | 17.65 | 700 | 0.16 | 2 | 50.00 | 246 | 0.70 | 1 | 100.00 | 185 | 0.54 |
| Feces (pooled) | 16 | 26.67 | 982 | 6.50 | 1 | 100.00 | 871 | 5.28 | 0 | - | 0 | - |
| Intestine | 13 | 0 | 65 | 0 | 0 | - | 0 | - | 0 | - | 0 | - |
| Kidney | 35 | 11.43 | 447 | 0.94 | 6 | 50.00 | 216 | 4.27 | 0 | - | 0 | - |
| Liver | 32 | 3.12 | 122 | 0.08 | 0 | - | 0 | - | 0 | - | 0 | - |
| Lung | 15 | 0 | 71 | 0 | 0 | - | 0 | - | 0 | - | 0 | - |
| Nasal | 5 | 60.00 | 1107 | 0.44 | 5 | 60.00 | 1107 | 0.44 | 0 | - | 0 | - |
| Oral | 128 | 16.41 | 11408 | 0.78 | 89 | 16.85 | 5928 | 1.09 | 7 | 71.43 | 3570 | 0.42 |
| Pooled swabs/ samples | 88 | 32.18 | 5344 | 5.40 | 40 | 25.00 | 2599 | 2.06 | 17 | 100 | 2299 | 22.54 |
| Pooled tissue | 69 | 31.88 | 2993 | 6.53 | 37 | 45.95 | 1278 | 9.70 | 7 | 71.43 | 1612 | 16.73 |
| Reproductive | 1 | 100 | 871 | 2.07 | 1 | 100 | 871 | 2.07 | 0 | - | 0 | - |
| Skin swab | 2 | 0 | 38 | 0 | 2 | 0 | 38 | 0 | 0 | - | 0 | - |
| Spleen | 15 | 71.43 | 1093 | 6.89 | 3 | 100 | 351 | 6.18 | 3 | 100 | 345 | 4.95 |
| Urinary (single) | 125 | 24.19 | 16787 | 1.68 | 105 | 20.95 | 10625 | 1.77 | 8 | 42.86 | 4244 | 0.10 |
| Urinary (pooled) | 315 | 38.43 | 45411 | 4.21 | 272 | 31.60 | 39449 | 2.06 | 4 | 100.00 | 1258 | 0.62 |
| Unspecified | 2 | 0 | 956 | 0 | 2 | 0 | 956 | 0 | 0 | - | 0 | - |
| **Viral Isolation** | | | | | | | | | | | | |
| Blood or serum | 2 | 0 | 277 | 0 | 2 | 0 | 277 | 0 | 0 | - | 0 | - |
| Brain | 5 | 0 | 200 | 0 | 1 | 0 | 5 | 0 | 2 | 0 | 159 | 0 |
| Fecal, rectal, anal | 7 | 0 | 176 | 0 | 4 | 0 | 108 | 0 | 0 | - | 0 | - |
| Feces (single) | 2 | 0 | 158 | 0 | 0 | - | 0 | - | 2 | 0 | 158 | 0 |
| Feces (pooled) | 1 | 0 | 4 | 0 | 0 | - | 0 | - | 1 | 0 | 4 | 0 |
| Heart | 3 | 0 | 167 | 0 | 1 | 0 | 5 | 0 | 2 | 0 | 162 | 0 |
| Intestine | 3 | 0 | 135 | 0 | 1 | 0 | 6 | 0 | 2 | 0 | 129 | 0 |
| Kidney | 10 | 10.00 | 479 | 0.95 | 4 | 0 | 207 | 0 | 4 | 25.00 | 225 | 2.38 |
| Liver | 6 | 0 | 248 | 0 | 4 | 0 | 207 | 0 | 2 | 0 | 41 | 0 |
| Lung | 7 | 0 | 417 | 0 | 4 | 0 | 207 | 0 | 3 | 0 | 210 | 0 |
| Oral | 14 | 0 | 1059 | 0 | 8 | 0 | 664 | 0 | 0 | - | 0 | - |
| Pooled tissue | 8 | 25.00 | 932 | 0.65 | 2 | 50.00 | 805 | 0.23 | 2 | 50.00 | 46 | 2.38 |
| Reproductive | 3 | 0 | 177 | 0 | 1 | 0 | 0 | 1.03 | 2 | 0 | 172 | 0 |
| Spleen | 7 | 0 | 264 | 0 | 2 | 0 | 11 | 0 | 3 | 0 | 201 | 0 |
| Urinary (single) | 10 | 20.00 | 733 | 0.07 | 6 | 33.33 | 652 | 0.12 | 0 | - | 0 | - |
| Urinary (pooled) | 21 | 94.74 | 2054 | 4.55 | 6 | 100.00 | 1062 | 4.78 | 13 | 100.00 | 511 | 4.09 |
| Urinary (unspecified) | 1 | 0 | 118 | 0 | 0 | - | 0 | - | 0 | - | 0 | - |
| Partially-eaten fruit | 1 | 100 | 27 | 3.70 | 1 | 100.00 | 27 | 3.70 | 0 | - | 0 | - |
