## Supplementary material for "Paramyxoviruses in Old World fruit bats (Pteropodidae): an open database and synthesis of sampling effort, viral positivity, and coevolution": Table S2

**Supplementary Table 2. Database breakdown by type of paramyxovirus.**
Number of studies, host species, and samples tested for different paramyxoviruses, with positivity rates reported for serological and PCR detection attempts as well as number of successful viral isolations.

| Virus | # studies | # bat species tested | # serological samples tested | Overall seropositivity (%) | # PCR samples tested | Overall PCR positivity (%) | # viral isolations |
|---|---|---|---|---|---|---|---|
| **Orthoparamyxovirinae** | | | | | | | |
| *Henipavirus* | | | | | | | |
| Angavokely virus | 1 | 1 | 0 | n/a | 96 | 12.4 | 0 |
| Cedar virus | 6 | 5 | 1267 | 8.4 | 872 | 0.2 | 3 |
| Ghanaian bat henipavirus | 1 | 1 | 0 | n/a | 215 | 1.4 | 0 |
| Hendra virus | 31 | 25 | 6570 | 17.3 | 43944 | 1.6 | 15 |
| Nipah virus | 56 | 34 | 19831 | 30.0 | 21882 | 1.8 | 7 |
| Other/unspecified henipaviruses | 7 | 15 | 810 | 34.9 | 1874 | 4.5 | 0 |
| "Henipa-like" orthoparamyxoviruses | 5 | 3 | 0 | n/a | 2806 | 14.3 | 0 |
| **Rubulavirinae** | | | | | | | |
| *Orthorubulavirus* | | | | | | | |
| Alston virus | 1 | 3 | 120 | 8.9 | 0 | n/a | 1 |
| Dawn bat paramyxovirus | 1 | 1 | 0 | n/a | 206 | 44.8 | 0 |
| Human mumps virus | 1 | 5 | 52 | (not reported) | 0 | n/a | 0 |
| *Pararubulavirus* | | | | | | | |
| Achimota viruses 1, 2, 3 | 3 | 1 | 946 | 17.7 | 0 | n/a | 3 |
| Hervey virus | 1 | Unspecified *Pteropus* | 0 | n/a | 0 | n/a | 9 |
| Menangle virus | 8 | 7 | 936 | 29.1 | 0 | n/a | 12 |
| Sosuga virus | 1 | 3 | 0 | n/a | 1611 | 2.9 | 0 |
| Teviot virus | 3 | 3 | 0 | n/a | 2003 | 0.5 | 17 |
| Tioman virus | 9 | 12 | 984 | 13.8 | 1 | 100.0 | 15 |
| Tuhoko viruses 1, 2, 3 | 1 | 1 | 69 | 60.9 | 996 | 1.5 | 0 |
| **Unclassified paramyxoviruses** | | | | | | | |
| Bat parainfluenza virus | 1 | 2 | 90 | 3.6 | 0 | n/a | 1 |
| Geelong paramyxovirus | 1 | 1 | 0 | n/a | 872 | 0.8 | 0 |
| Grove virus | 1 | Unspecified *Pteropus* | 0 | n/a | 0 | n/a | 10 |
| Yarra bend paramyxovirus | 2 | 3 | 0 | n/a | 1874 | 2.9 | 0 |
| Yeppoon virus | 1 | Unspecified *Pteropus* | 0 | n/a | 0 | n/a | 8 |
