## Supplementary material for "Paramyxoviruses in Old World fruit bats (Pteropodidae): an open database and synthesis of sampling effort, viral positivity, and coevolution": Table S3

**Supplementary Table 3. Database breakdown by country.**
Number of studies, number of samples, and mean viral prevalence and seropositivity rates for all countries.

| Country | UN Geoscheme region | Number of studies | Number of samples | Average PCR prevalence (%) | Average seroprevalence (%) |
|---|---|---|---|---|---|
| Australia | Oceania | 29 | 58590 | 4.02 | 29.99 |
| Bangladesh | Asia | 8 | 22885 | 1.95 | 20.74 |
| Cambodia | Asia | 3 | 5356 | 0.62 | 3.73 |
| China | Asia | 5 | 1301 | 1.35 | 35.64 |
| Comoros | Africa | 1 | 26 | 0 | NA |
| Democratic Republic of the Congo | Africa | 2 | 146 | 6.08 | NA |
| Equatorial Guinea | Africa | 5 | 954 | 0 | 37.88 |
| Fiji | Oceania | 1 | 68 | 0 | NA |
| Gabon | Africa | 1 | 21 | 0 | NA |
| Ghana | Africa | 13 | 8189 | 11.23 | 44.88 |
| India | Asia | 13 | 3970 | 3.05 | 19.17 |
| Indonesia | Asia | 6 | 1407 | 1.37 | 22.31 |
| Kenya | Africa | 3 | 521 | 1.74 | NA |
| Madagascar | Africa | 5 | 4114 | 1.74 | 10.85 |
| Malawi | Africa | 3 | 38 | 0 | 31.25 |
| Malaysia | Asia | 9 | 5396 | 1.19 | 16.42 |
| Mayotte | Africa | 1 | 44 | 0 | NA |
| Myanmar | Asia | 1 | 58 | 0 | NA |
| Nigeria | Africa | 2 | 87 | 0 | NA |
| Papua New Guinea | Oceania | 3 | 1660 | 0 | 16.65 |
| Republic of Congo | Africa | 2 | 329 | 4.77 | NA |
| Rwanda | Africa | 1 | 28 | 6.25 | NA |
| Sao Tome and Principe | Africa | 4 | 566 | NA | 32.25 |
| Saudi Arabia | Asia | 1 | 26 | 0 | NA |
| Singapore | Asia | 3 | 360 | 24.52 | NA |
| South Africa | Africa | 3 | 3333 | 4.38 | NA |
| Tanzania | Africa | 4 | 825 | 20 | 29.71 |
| Thailand | Asia | 6 | 7353 | 1.64 | 5.91 |
| Timor-Leste | Asia | 1 | 734 | 0.78 | 11.54 |
| Uganda | Africa | 4 | 1626 | 16.73 | 96.43 |
| Vietnam | Asia | 2 | 329 | 4.85 | 22.32 |
| Zambia | Africa | 4 | 341 | 7.89 | 42.92 |
