## Supplementary material for "Paramyxoviruses in Old World fruit bats (Pteropodidae): an open database and synthesis of sampling effort, viral positivity, and coevolution": Table S4

**Supplementary Table 4. Results from PACo cophylogenetic analysis of paramyxoviruses in Pteropodidae.**

Jackknifed squared residuals and upper 95% confidence intervals for each known pteropodid–paramyxovirus link from PACo.

| Host | Virus | Mean jackknifed squared residual | Upper 95% confidence limit |
|---|---|---|---|
| *Pteropus vampyrus* | *Tioman pararubulavirus* | 432.89 | 884.50 |
| *Pteropus giganteus* | *Tioman pararubulavirus* | 431.80 | 884.51 |
| *Pteropus hypomelanus* | *Tioman pararubulavirus* | 473.38 | 920.89 |
| *Pteropus poliocephalus* | *Teviot pararubulavirus* | 491.25 | 933.12 |
| *Pteropus alecto* | *Menangle pararubulavirus* | 422.72 | 863.00 |
| *Rousettus leschenaultii* | *Tuhoko pararubulavirus 1* | 655.50 | 1158.16 |
| *Eidolon helvum* | *Achimota pararubulavirus 2* | 905.47 | 1318.82 |
| *Rousettus leschenaultii* | *Tuhoko pararubulavirus 2* | 636.65 | 1160.12 |
| *Eidolon helvum* | *Achimota pararubulavirus 1* | 912.95 | 1328.74 |
| *Rousettus leschenaultii* | *Tuhoko pararubulavirus 3* | 584.58 | 1112.79 |
| *Rousettus aegyptiacus* | *Sosuga pararubulavirus* | 572.53 | 1106.07 |
| *Pteropus scapulatus* | *Hendra henipavirus* | 108.61 | 263.90 |
| *Pteropus poliocephalus* | *Hendra henipavirus* | 90.05 | 239.64 |
| *Pteropus conspicillatus* | *Hendra henipavirus* | 100.07 | 260.27 |
| *Pteropus alecto* | *Hendra henipavirus* | 99.63 | 260.27 |
| *Rousettus amplexicaudatus* | *Nipah henipavirus* | 1260.18 | 1776.44 |
| *Rousettus leschenaultii* | *Nipah henipavirus* | 1399.99 | 1873.03 |
| *Pteropus vampyrus* | *Nipah henipavirus* | 102.63 | 260.90 |
| *Pteropus giganteus* | *Nipah henipavirus* | 102.70 | 266.71 |
| *Pteropus lylei* | *Nipah henipavirus* | 103.01 | 268.49 |
| *Pteropus hypomelanus* | *Nipah henipavirus* | 99.54 | 256.13 |
| *Pteropus poliocephalus* | *Cedar henipavirus* | 69.96 | 221.75 |
| *Eidolon helvum* | *Ghana henipavirus* | 1133.24 | 1396.15 |
