## Supplementary figures and images for "Paramyxoviruses in Old World fruit bats (Pteropodidae): an open database and synthesis of sampling effort, viral positivity, and coevolution"

### Fig. S1

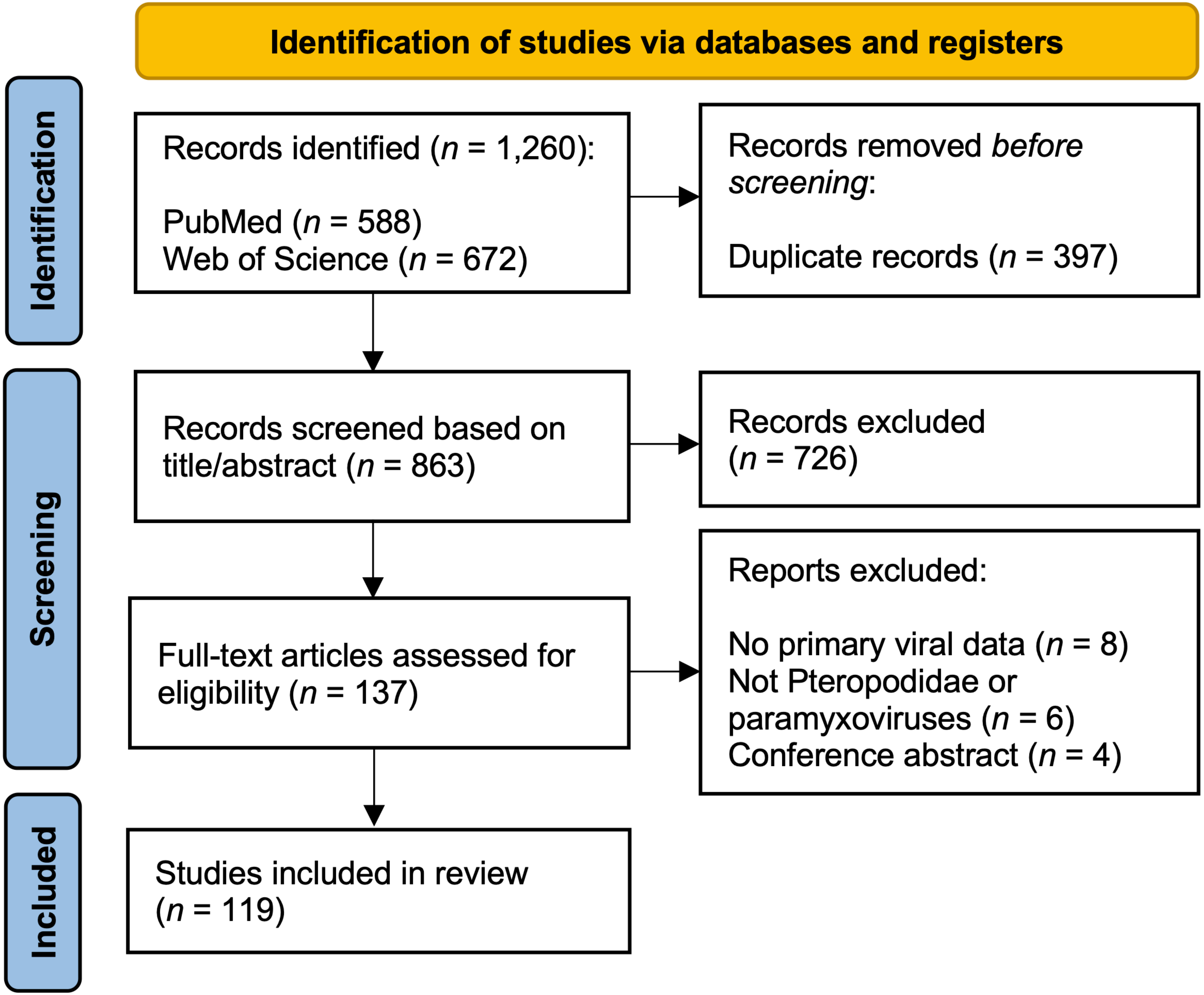
